# Hypertension-mediated cardiac fibrosis is a mechanical process initiated by smooth muscle cells

**DOI:** 10.64898/2026.05.28.728606

**Authors:** Anna Kaganovsky, Anas Odeh, Shelly Eilot, Lavi Coren, Maher Abu-Saleh, Ariel Shemesh, Rotem Bela Shimron, Rohtem Aviram, Haguy Wolfenson, Izhak Kehat, Peleg Hasson

## Abstract

Hypertension represents the most prevalent chronic cardiovascular condition, typically culminating in pathological cardiac remodeling characterized by hypertrophy and extensive fibrosis. Although the cellular phenotypes associated with these changes are well-documented, the precise mechanisms by which hypertensive stress is sensed and transduced into a fibrotic program remain poorly defined. To elucidate these mechanisms, we investigated the role of Lysyl oxidase (LOX), an extracellular matrix (ECM)-modifying enzyme that is upregulated during hypertensive stress and associated with cardiovascular diseases. By employing cell-type specific Cre-Lox technology to conditionally delete *Lysyl oxidase* in either smooth muscle cells (SMCs) or fibroblasts, the primary ECM-secreting cell populations, we demonstrate that fibroblast-specific *Lox* deletion had no significant impact on the progression of cardiac fibrosis. Conversely, SMC-specific *Lox* deletion selectively inhibited the fibrotic response without affecting other remodeling parameters, such as cardiac hypertrophy. Notably, in the SMC-specific *Lox* knockout hearts, fibrosis was restricted to the perivascular niche and failed to propagate into the cardiac interstitium. We find that this transition is a mechanical, ECM-dependent process initiated by SMCs. Our results identify SMCs, rather than fibroblasts, as the primary sensors and initiators of the hypertensive fibrotic response. These findings demonstrate that fibrosis can be uncoupled from other hypertensive manifestations and identify SMC-mediated ECM modification as a potential therapeutic target for treating hypertensive heart disease.

## Introduction

The cardiovascular system senses and adapts to distinct stressful conditions. While these responses enable cardiovascular function under demanding circumstances, upon prolonged stress these adaptations often promote cardiac pathologies. Hypertension, the persistently elevated arterial blood pressure, is the most common chronic stressful condition^1^. The hypertension-dependent cardiac remodeling events are varied and depend on the specific cell type^2^. While cardiomyocytes grow in size, increasing sarcomere number, fibroblasts and smooth muscle cells (SMC) increase the secretion of extracellular matrix (ECM). However, often these remodeling processes are pathological leading to reduced cardiac relaxation and contractile dysfunction. In such cases, cardiac fibroblasts differentiate into myofibroblasts, promoting excessive ECM secretion and fibrotic tissue accumulation, often at the expense of resident cardiac cells^3^. Notably, these scarred areas alter the cellular microenviroment, lead to adverse mechanical signals that impair contractility and relaxation dynamics, ultimately leading to cardiac failure^4–7^. Although these events have been highly documented, how hypertension induces them and which cells initiate the response remain poorly understood.

The Renin-angiotensin-aldosterone system (RAAS) is a major regulator of blood pressure and its overactivation is key to human hypertension. Accordingly, it is associated with heart failure. Angiotensin II (AngII), is the principle active hormone generated by the RAAS system and the primary effector molecule that directly regulates blood pressure and as such AngII infusion is the most widely used method for experimentally inducing hypertension in rodents^8^. Although inducing hypertension, AngII also promotes cardiac fibrosis through AT₁-receptor signaling in multiple cardiac cell types. In SMCs, AngII induces strong vasoconstriction, raising afterload and mechanical stress on the myocardium. In cardiomyocytes, AngII activates hypertrophic signaling pathways and increases oxidative stress, which further elevates wall tension and stimulates profibrotic cytokine release. At the same time, AngII directly drives cardiac fibroblast proliferation and myofibroblast differentiation through TGF-β1, reactive oxygen species (ROS), and MAPK pathways, leading to excessive collagen deposition and reduced ECM degradation. Together, these mechanical and biochemical effects create a feed-forward loop that accelerates fibrosis and myocardial stiffening.

The augmented ECM deposition and the ensuing fibrosis in response to the increased cardiac workload are also associated with a significant upregulation of ECM modifying enzymes. One such key enzyme is LYSYL OXIDASE (LOX), a secreted collagen and elastin crosslinking enzyme^9^. Recent work has demonstrated LOX and LOX-Like family members are also essential for initiating fibronectin (FN) fibrillogenesis inducing integrin activation^10,11^. These steps are necessary for the assembly of collagen and other ECM molecules, positioning lysyl oxidases as the initiators of ECM accumulation and the fibrotic response. LOX is highly expressed by multiple cell types within the cardiovascular system, including cardiomyocytes, although it is most highly expressed by SMC and fibroblasts. Accordingly, LOX activity is associated with cardiovascular development and diseases^12–14^. In response to hypertension, LOX expression is upregulated^15^, presumably to enable remodeling of the newly secreted ECM facilitating the ability of the aorta and cardiac muscle to withstand increased mechanical pressure.

Using Cre lines under the regulation of *platelet-derived growth factor receptor α* (*PDGFRα*) and *Myosin heavy chain 11 (Myh11)*^16–18^ expressed in fibroblasts and SMC, respectively, we set to delete Lox in either cell type in hypertensive mice. While much focus has been placed on the roles of fibroblasts in regulating the fibrotic reaction, our results highlight an unexpected role for the SMC in the process. Surprisingly, we find that SMCs, rather than fibroblasts, are the cells that sense and induce the fibrotic response following hypertension. Specifically, we demonstrate that in response to hypertension, SMC-remodeled ECM initiates fibroblast activation linking these two processes. We demonstrate that deleting Lox specifically in the SMC, but not in fibroblasts, uncouples the two highly associated processes, hypertension and cardiac fibrosis. These findings open avenues for novel interventions into the adverse effects of this common pathological condition.

## Results

### SMC-derived LOX is essential for cardiac remodeling

The cardiovascular system is the primary tissue that adapts to increased cardiac workload, such as that caused by hypertension, through induction of mechanical pressure by vascular cells. The cardiac remodeling process is highly associated with increased ECM deposition and fibrosis. To dissect the processes underlying its development, we focused on Lox, a key ECM remodeling enzyme which has been demonstrated to be involved in the regulation of these processes within the cardiovascular system^19^. To identify the cells that express *Lox*, we followed its expression using fluorescent RNA in situ hybridization (FISH). We find that *Lox* is primarily expressed by fibroblasts and SMC, the principal cells secreting cardiac ECM (Fig. 1A-A’’). The *Lox* expression in the ECM secreting cells, led us to examine whether hypertension affects its expression. Towards that end, at the age of 3 months, osmotic minipumps (Alzet) infused with 10mg/kg/day AngII were subcutaneously implanted for 28 days and the mice were then harvested at 4 months of age (Fig. 1B). Western blot analysis of baseline and hypertensive hearts demonstrates Lox is significantly upregulated following AngII-mediated hypertension (Fig 1C,D).

**Figure 1.**
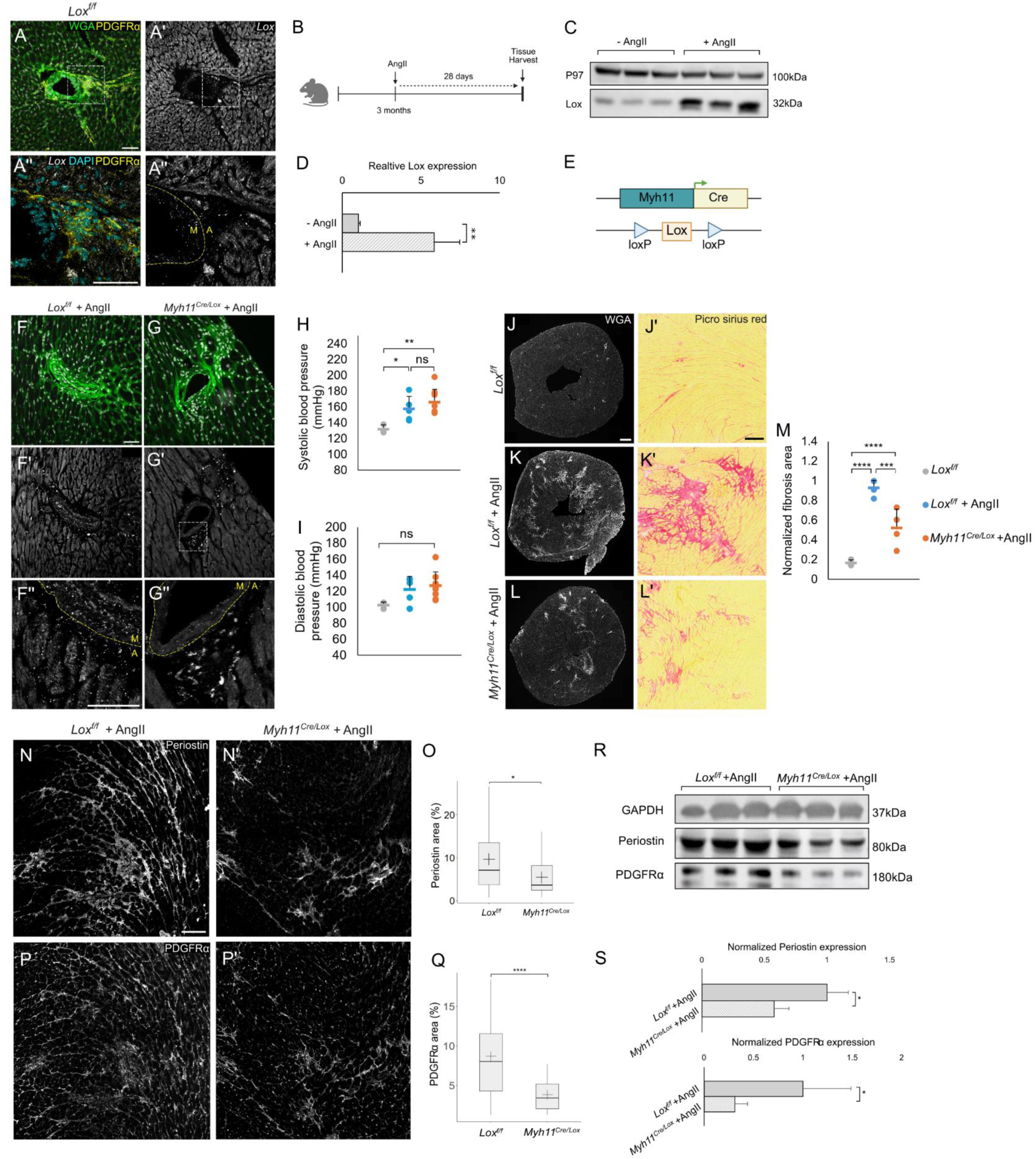
*Lox* in SMC attenuates hypertension-mediated fibrosis. FISH for *Lox* (grey) in baseline *Lox^f/f^* mice demonstrates its expression in vascular media (M) and adventitia (A) marked by PDGFRα (yellow). WGA (green) highlights cell membranes and general ECM (A) (Scale bars 50µm). Experimental scheme (B). Western blot (C) for Lox and quantification (D) from control (-AngII) (N=3) and AngII-treated (N=3) mice. Schematic illustration of the mouse model used for SMC-specific *Lox* deletion (E). FISH for *Lox* in control *Lox^f/f^* (F) and *Myh11^Cre/Lox^* (G) hypertensive mice, validating *Lox* deletion in SMC (Scale bars 50µm). Systolic (H) and diastolic (I) blood pressure measurements of baseline *Lox^f/f^* (N=3) and hypertensive AngII-treated *Lox^f/f^* (N=5) and *Myh11^Cre/Lox^* (N=8) mice. WGA and Sirius red staining (J-L) (Scale bars 500µm, 200µm) and quantification (M) of baseline *Lox^f/f^* (J) (N=3), hypertensive AngII-treated *Lox^f/f^* (N=6)(K) and *Myh11^Cre/Lox^* (N=4)(L) mice. Immunostaining for Periostin (N) (Scale bar 100µm) and quantification (O) (n=31, *Lox^f/f^* and n=19 *Myh11^Cre/Lox^* mice) and PDGFRα (P,Q) (n=47, *Lox^f/f^* and n=31 *Myh11^Cre/Lox^* mice) in hypertensive AngII-treated *Lox^f/f^* (N=6) and *Myh11^Cre/Lox^* (N=4) mice. Western blot (R) and quantification (S) for Periostin and PDGFRα in hypertensive AngII-treated *Lox^f/f^* (N=3) and *Myh11^Cre/Lox^* (N=3) mice. In A’’, F’’ and G’’ A designates adventitia and M designates media. Data are presented as the mean ± SD. Statistical significance is represented by an asterisk as calculated using one-way ANOVA and post hoc Tukey’s test and Unpaired t-test. n.s. = not significant. *p < 0.05;**p < 0.01; ***p < 0.001.; ****p < 0.0001).

Having seen that Lox is upregulated in AngII-infused hypertensive hearts, we investigated whether it plays a role in hypertension-mediated cardiac remodeling. Towards that end, we took advantage of the Cre line driven by the SMC-specific *Myh11* promoter (*Myh11^Cre^*; JAX #7742)^16^ and crossed it to the *Lox* conditional allele (*Lox^f/f^*)^20^ (Fig. 1E). Crossing this Cre line to a *Rosa26nTnG* reporter (JAX #023537^21^) demonstrated the Cre’s specificity to SMCs (Fig. S1A). FISH analysis verified this Cre line specifically deletes Lox in SMC (Fig. 1F,G). As recently demonstrated, in otherwise homeostatic *Myh11^Cre^; Lox^f/f^* (*Myh11^Cre/Lox^*) mice, no gross aortic media organization or structure, nor blood pressure changes were observed following this deletion even though western blot analysis demonstrated Lox expression was reduced by 60-80% in mutant aortas^22^.

Hypertension induction was carried out as described above in 3 months old *Lox^f/f^* as control, and *Myh11^Cre/Lox^* mice. Blood pressure was monitored as early as 5 days following AngII administration. Even though hypertensive *Lox* mutant mice develop aneurysms^22^, similar hypertension levels were observed in control and *Myh11^Cre/Lox^* mice suggesting that even in the lack of *Lox*, SMC still respond to AngII (Fig. 1H,I). Strikingly, monitoring ECM deposition using wheat germ agglutinin (WGA) and Sirius red stainings, highlighted a substantial difference between the two mouse lines where in *Myh11^Cre/Lox^*mice, a significantly reduced fibrotic accumulation was observed (Fig. 1J-M). Immunostaining and western blots for PDGFRα, a marker of fibroblasts, and Periostin (Postn), a marker of activated fibroblasts^23^, demonstrated a similar trend where in hypertensive mice lacking Lox in the SMC, significantly less fibroblasts and activated fibroblasts were observed (Fig. 1N-S).

To further strengthen these observations and exclude the possibility they resulted from early embryonic *Lox* deletion in cells that had transiently expressed *Myh11*, we utilized an SMC-specific tamoxifen-inducible *Myh11^Cre^* line (*Myh11^CreERT^*^2^; JAX#019079)^18^ to induce *Lox* deletion at later stages. Tamoxifen was administered for five consecutive days starting at 3 weeks of age, followed by monthly maintenance doses. Similar to the non-inducible Cre model described above, Cre activity was restricted to the medial layer of the cardiac vessels (Fig. S2A). At 3 months of age, mice were implanted with AngII-infused minipumps and harvested one month later. Notably, this inducible deletion strategy also resulted in a significant reduction in cardiac fibrosis, supporting the conclusion that the phenotype is specifically driven by *Lox* deletion in SMCs (Fig. S2B-D).

Apart from increased ECM deposition, the cardiac remodeling response to hypertension is a complex, multistep process involving cardiac hypertrophy affecting the vasculature ultimately leading to impaired cardiac function. To dissect whether the reduced Lox in SMC affected all cardiac remodeling processes or whether it specifically affected the fibrotic response, we assessed several parameters associated with the hypertension stress response. As expected, hypertensive control mice subjected to prolonged AngII treatment exhibited hallmark features of cardiac remodeling, including increased cardiomyocyte cross-sectional area (CSA) (Fig. 2A-H) and heartweight to body weight (HW/BW) ratio (Fig. 2I). No differences in these hypetrophy-associated parameters were observed between control and *Myh11^Cre/Lox^* hypertensive mice (Fig. 2A-I). Further, no significant changes to cardiac function, as measured by fractional shortening (FS) were noted (Fig. 2J). Quantification of the total number of blood vessels to heart tissue area demonstrated that also in this parameter no differences were observed (Fig. 2K). however, while the area of small arterioles (marked by αSMA expression), was not altered between the two mouse lines, a significant increase in the area of medium (100-2000 µm^2^) sized vessels was observed in the *Myh11^Cre/Lox^* mice (Fig. 2L,M) suggesting the vessels in the *Myh11^Cre/Lox^* hypertensive mice are somewhat larger. Altogether, these findings indicate that smooth muscle cell–derived Lox is critical for promoting the fibrosis-associated cardiac remodeling process in response to hypertension. Importantly, these results further demonstrate that hypertension and cardiac fibrosis, two processes considered to be highly associated, can be functionally uncoupled.

**Figure 2.**
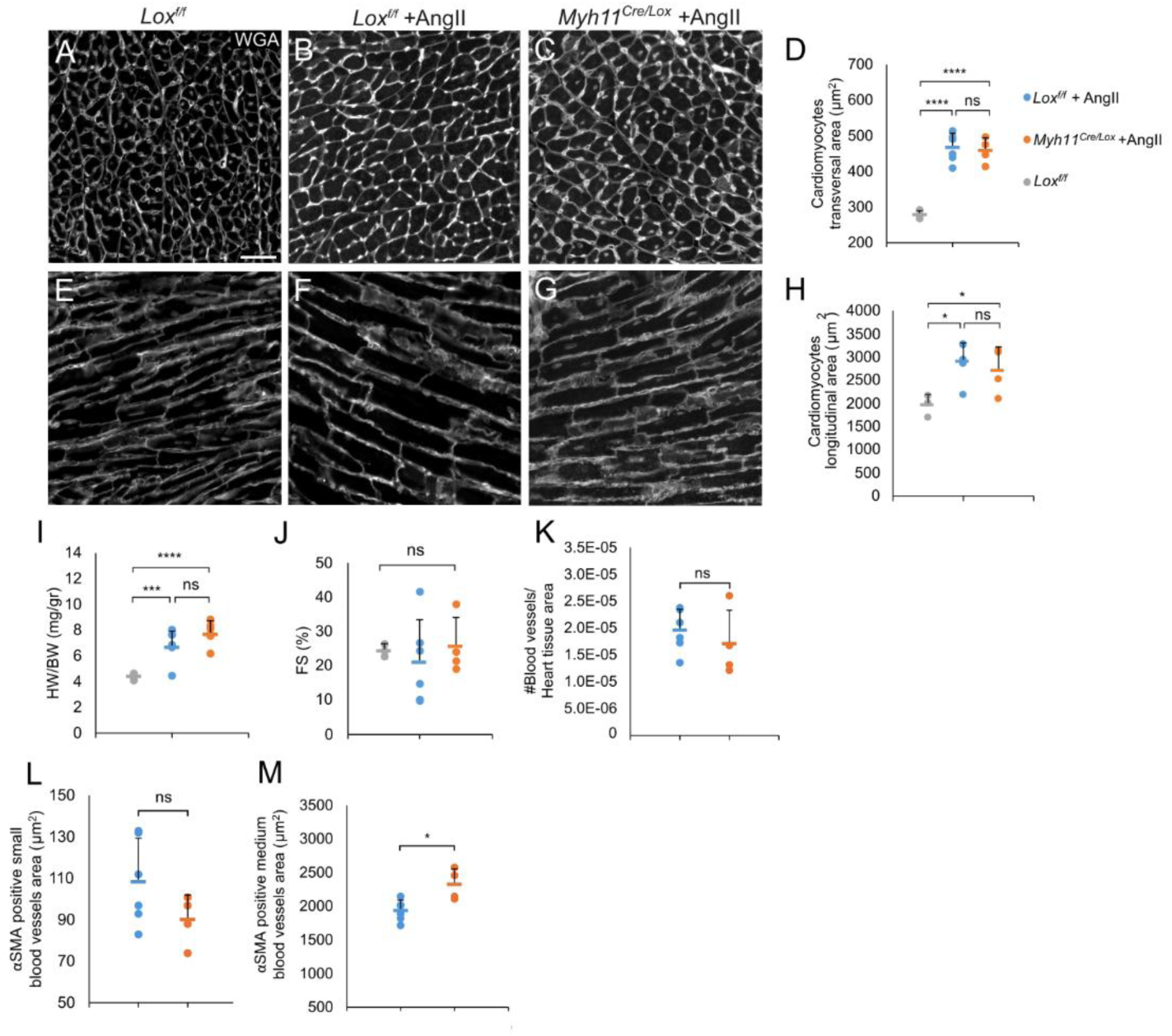
*Lox* deletion in SMC does not affect cardiac hypertrophy. WGA staining (A-C, E-G)(Scale bar 50µm) and quantifications of transverse (A-C) and longitudinal (E-G) axes of baseline *Lox^f/f^* (N=3),AngII-treated *Lox^f/f^* (N=6) and *Myh11^Cre/Lox^* (N=3) mice (D,H). HW/BW ratios of baseline (N=3) and AngII-treated *Lox^f/f^* (N=8) and *Myh11^Cre/Lox^* (N=7) mice (I). FS(%) marking cardiac function of baseline *Lox^f/f^* (N=3), AngII-treated *Lox^f/f^* (N=6) and *Myh11^Cre/Lox^* (N=4) mice (J). Quantification of total number of αSMA-positive blood vessels/ heart tissue area (K), of αSMA-positive blood small (0-100µm) (L) and medium (100-2000µm) (M) sized vasculature of baseline *Lox^f/f^* (N=3), AngII-treated *Lox^f/f^* (N=6) and *Myh11^Cre/Lox^* (N=4) mice. Data are presented as the mean ± SD. Statistical significance is represented by an asterisk as calculated using one-way ANOVA and post hoc Tukey’s test and Unpaired t-test. n.s. = not significant. *p < 0.05;***p < 0.001.; ****p < 0.0001.

### Lox expression in fibroblasts is not essential for inducing cardiac remodeling

Modifications to the ECM are a hallmark of tissue remodeling processes primarily mediated by fibroblasts and myofibroblasts^3^. The majority of myofibroblasts arise from resident fibroblasts, with a subset derived from perivascular cells^24^ and other sources^3^. Hence, the surprising observations that following Lox deletion specifically in SMC, cardiac fibrosis was attenuated in hypertensive hearts prompted us to test whether this phenotype is attributable to (i) SMC-derived myofibroblasts or (ii) unintended, low-level *Myh11^Cre^* activity in fibroblasts. To directly test these possibilities, we set to delete *Lox* specifically in fibroblasts, cells which express high Lox levels under normal conditions and upregulate it further in response to hypertension (Fig. 1) or other cardiac stresses^25,26^. To achieve fibroblast-specific deletion, we took advantage of the *PDGFRα^CreERT^*^2^ mice, a Tamoxifen inducible Cre line driven by the *PDGFRα* promoter, highly expressed in fibroblasts^17^. Reporter gene activation (*PDGFRα^CreERT^*^2^; *Rosa26nTnG*) demonstrates its robust specific activation following TM administration (Fig. S3).

Upon weaning, at the age of 3-4 weeks, *PDGFRα^CreERT^*^2^*; Lox^f/f^* (Fig. 3A) and *control* (*Lox^f/f^* or *PDGFRα^CreERT^*^2^*^/Lox^*) were intra-peritonealy (IP) injected with TM (100µl, 20mg/ml) for 5 consecutive days and then on once a month. At 3 months of age, mice were implanted with AngII-infused osmotic minipumps as described above (Fig. 3B). FISH analyses verified the depletion of *Lox* mRNA in fibroblasts (Fig. 3C,D). Blood pressure measurements confirmed hypertension induction. Notably, as with *Lox* deletion in SMC, no differences were observed between the control group and *PDGFRα^CreERT^*^2^*^/Lox^*mice (Fig. 3E,F) demonstrating that this *Lox* deletion regime does not affect SMC ability to induce hypertension. One month later, at 4 months of age, mice were subjected to echocardiography to monitor cardiac function and were then harvested.

**Figure 3.**
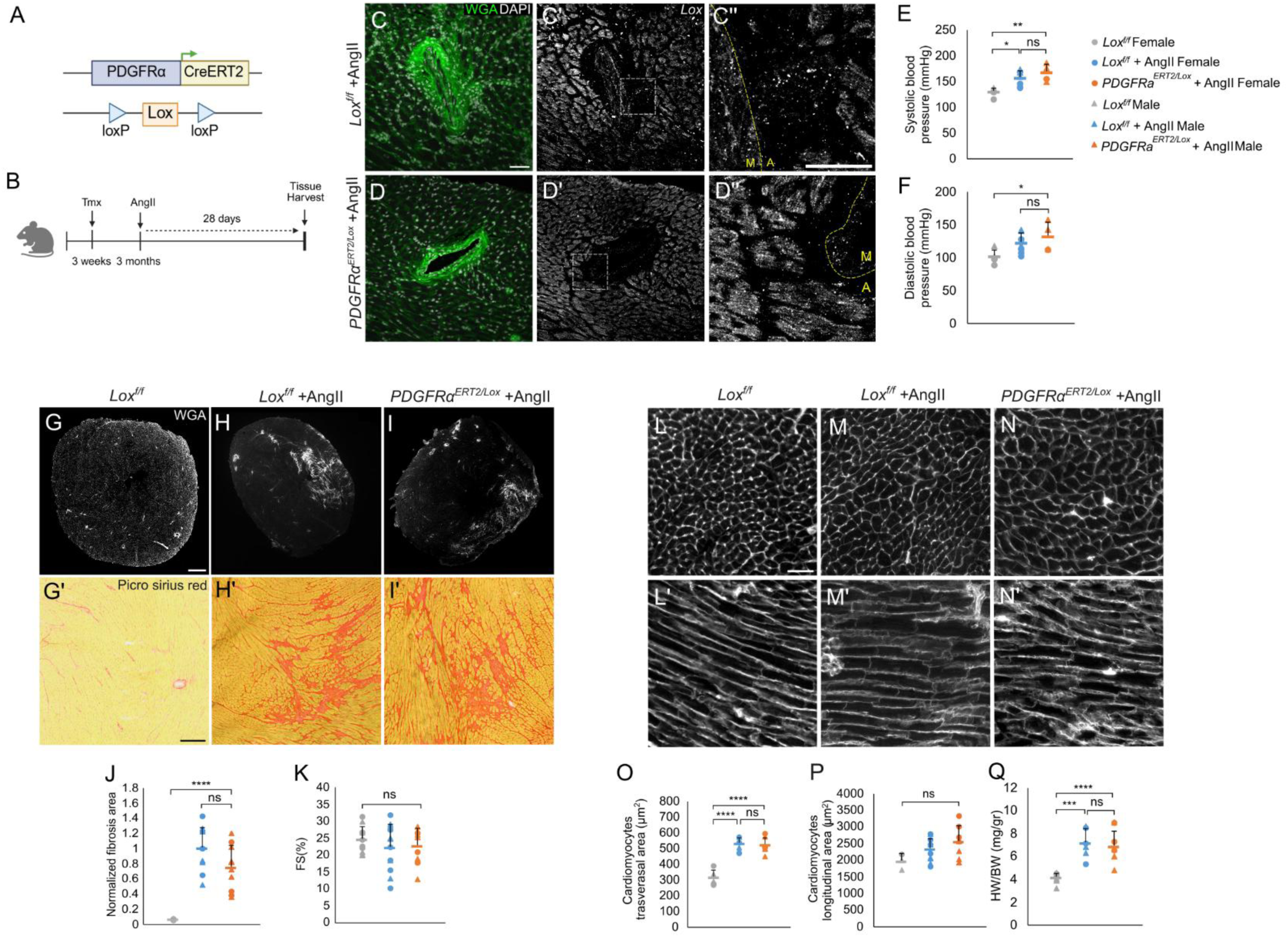
*Lox* deletion in fibroblasts does not affect hypertension-mediated cardiac remodeling. Schematic illustration of the mouse model used for fibroblast-specific *Lox* deletion (A) and experimental scheme (B). FISH for *Lox* in hypertensive control *Lox^f/f^* (C) and *PDGFRα^ERT^*^2^*^/Lox^* (D) mice, demonstrating *Lox* expression is significantly downregulated in fibroblasts. WGA (green) marking cell membranes and ECM (Scale bar 50µm). In D’’, A designates adventitia and M media. Systolic (E) and diastolic (F) blood pressure measurements of baseline *Lox^f/f^* (N=5), AngII-treated *Lox^f/f^* (N=7) and *Myh11^Cre/Lox^* (N=4) mice. WGA (top) (Scale bar 500µm) and Sirius red (bottom) (Scale bar 200µm) staining of baseline *Lox^f/f^* (G)(N=6), hypertensive AngII-treated *Lox^f/f^* (H)(N=11) and *PDGFRα^ERT^*^2^*^/Lox^* (I)(N=10) mice. Quantification of fibrotic area(%), highlighting no differences between the two hypertensive mouse lines (J). FS(%) demonstrates no changes to cardiac activity between baseline *Lox^f/f^* (N=5), hypertensive AngII-treated *Lox^f/f^* (N=12) and *PDGFRα^ERT^*^2^*^/Lox^* (N=8)(K). WGA staining (Scale bar 50µm) (L-N) and quantifications of cardiomyocytes’ CSA in transverse (O) and longitudinal (P) axes demonstrate similar hypertrophic responses in the hypertensive control *Lox^f/f^*(N=11) and *PDGFRα^ERT^*^2^*^/Lox^* (N=10) mice, relative to the baseline *Lox^f/f^* mice (N=3). HW/BW ratios (Q) demonstrate similar hypertrophic responses in the two hypertensive control *Lox^f/f^* (N=6) and *PDGFRα^ERT^*^2^*^/Lox^* (N=9) mice relative to the baseline *Lox^f/f^* mice (N=8). Data are presented as the mean ± SD. Statistical significance is represented by an asterisk as calculated using one-way ANOVA and post hoc Tukey’s test. n.s. = not significant. *p < 0.05;**p < 0.01; ***p < 0.001.; ****p < 0.0001.

Surprisingly, no differences in fibrosis accumulation marked by WGA and Sirius red staining (Fig. 3G-J) were observed between the two mouse lines. Along these lines, no other aspects of cardiac remodeling including reduced FS (Fig. 3K), increase in CSA and HW/BW were differentially regulated between the AngII-hypertensive control and *PDGFRα^CreERT^*^2^*^/Lox^*mice (Fig. 3L-Q). Altogether these results suggest that fibroblasts-derived Lox does not play a key role in the induction of the cardiac remodeling processes in response to hypertension.

### Smooth muscle cells shape the pattern of fibrotic propagation

In response to prolonged hypertension, fibrotic processes have been suggested to originate in the perivascular regions and progressively expand into the interstitial space^27^; however, the mechanisms governing this transition have remained poorly understood^28^. The reduced fibrosis accumulation in the *Myh11^Cre/Lox^* hypertensive mice (Fig. 1) led us to examine whether this phenotype reflects not only decreased ECM deposition but also differences in its spatial organization. As a first step, we monitored fibrotic spatial distribution. To investigate this, we quantified perivascular (Fig. 4A) and interstitial (Fig. 4B) collagen in the control and *Myh11^Cre/Lox^* hypertensive mice. To normalize the different vessel sizes, medial and adventitial collagen expression were divided by vessel area and this ratio was then marked as perivascular collagen. The remaining collagen, normalized to cardiac area, was considered as interstitial collagen, representing the fibrotic spread beyond the vascular niche. We find that the ratio of perivascular collagen to vessel area was similar between the control and *Myh11^Cre/Lox^* mouse lines’ hearts. In contrast, that of interstitial collagen was significantly reduced in the *Myh11^Cre/Lox^* (Fig. 4A-D).

**Figure 4.**
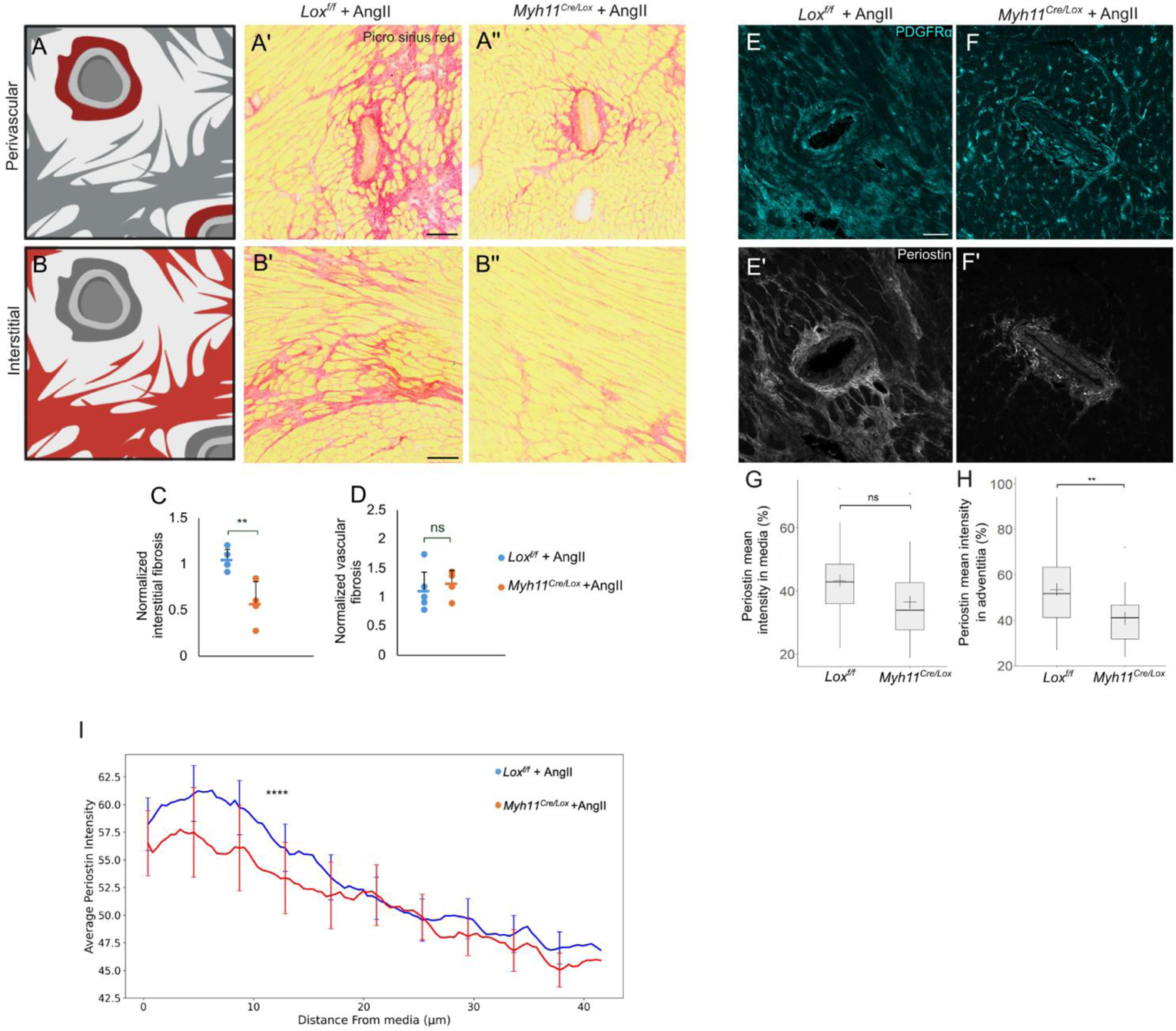
Fibrotic pattern is regulated by SMC-derived Lox. Schematic illustration of perivascular (A) and interstitial fibrosis (B). Sirius red staining of AngII-treated control *Lox^f/f^* (A’, B’) and *Myh11^Cre/Lox^* (A’’,B’’) mice hearts (Scale bar 100µm). Quantification of interstitial (C) and perivascular (D) fibrosis of hypertensive AngII-treated *Lox^f/f^* (N=6) and *Myh11^Cre/Lox^* (N=4) mice. Immunostaining for PDGFRα (cyan) and Periostin (grey) in AngII-treated *Lox^f/f^* (N=6, n=27) (E) and *Myh11^Cre/Lox^* (N=4, n=18) (F) mice hearts (Scale bar 50µm). Quantification of medial (G) and adventitial Periostin (H), highlighting reduced fibroblast activation in mutant mice adventitia. Line graph depicting average Postn intensity highlighting its distribution within the adventitia (I). Data are presented as the mean ± SD. Statistical significance is represented by an asterisk as calculated using Unpaired t-test (D,G) or KS test (H).n.s. = not significant. **p < 0.01, ****p<0.0001.

Having seen the reduced fibrotic scattering in the *Myh11^Cre/Lox^*mice, we wished to dissect the underlying cause. We therefore monitored the distribution of activated fibroblasts marked by Postn in the media and the surrounding vascular adventitia. Immunostaining for Postn together with PDGFRα (a fibroblast marker) shows that, Postn levels are similarly distributed in the media in both mouse lines. In contrast, fibroblast activation is significantly reduced in the adventitia of *Myh11^Cre/Lox^* hypertensive mice (Fig. 4E–I), suggesting that the effective range of the mechanical ‘stress signal’ is attenuated in mutant hearts. Together, these findings suggest that a smooth muscle cell–derived signal, mediated by Lox, facilitates fibroblast activation to drive the propagation of perivascular fibrosis into the cardiac interstitium.

### SMC-derived perivascular ECM organization regulates response to hypertension

In response to stressors such as hypertension, SMCs rapidly remodel their surrounding environment^29^. Cellular Fibronectin (cFN) is one of the earliest ECM core components to be affected, functioning both as a scaffold for collagen and other ECM proteins and as a mediator of mechanical signaling from the media to adjacent fibroblasts^30^. We recently demonstrated that lysyl oxidases (LOX and LOX-like 1–4) bind to and oxidize FN on domains shared by all FN isoforms. In turn, this oxidation affects FN fibrillogenesis and spatial organization^10,11^. We therefore examined whether vascular cFN distribution differs between control and *Myh11^Cre/Lox^* mice to determine whether the initial hypertensive response is altered in the mutant animals. Using a cFN-specific antibody at limiting dilutions we specifically marked its expression in the vascular media. We then segmented the region and applied deep learning–based texture analysis to extract Haralick features, which quantitatively describe image texture parameters such as dissimilarity, correlation, contrast, and homogeneity. In control vessels (N=6, n = 41), cFN expression displayed a consistent pattern corresponding to a narrow distribution peak suggesting its organization in the hypertensive media is a tightly regulated process. In contrast, mutant vessels (N = 4, n = 48) displayed disrupted cFN organization, characterized by a broad distribution of these parameters, indicative of increased spatial heterogeneity and loss of the tightly regulated fiber architecture observed in the control media. (Fig. 5A-C’’’). Similar disruptions in cFN expression were observed also in *Myh11^CreERT^*^2^*^/Lox^* further supporting the conclusion that these alterations are not a consequence of early embryonic defects in FN organization(Fig. S2E-F).

**Figure 5.**
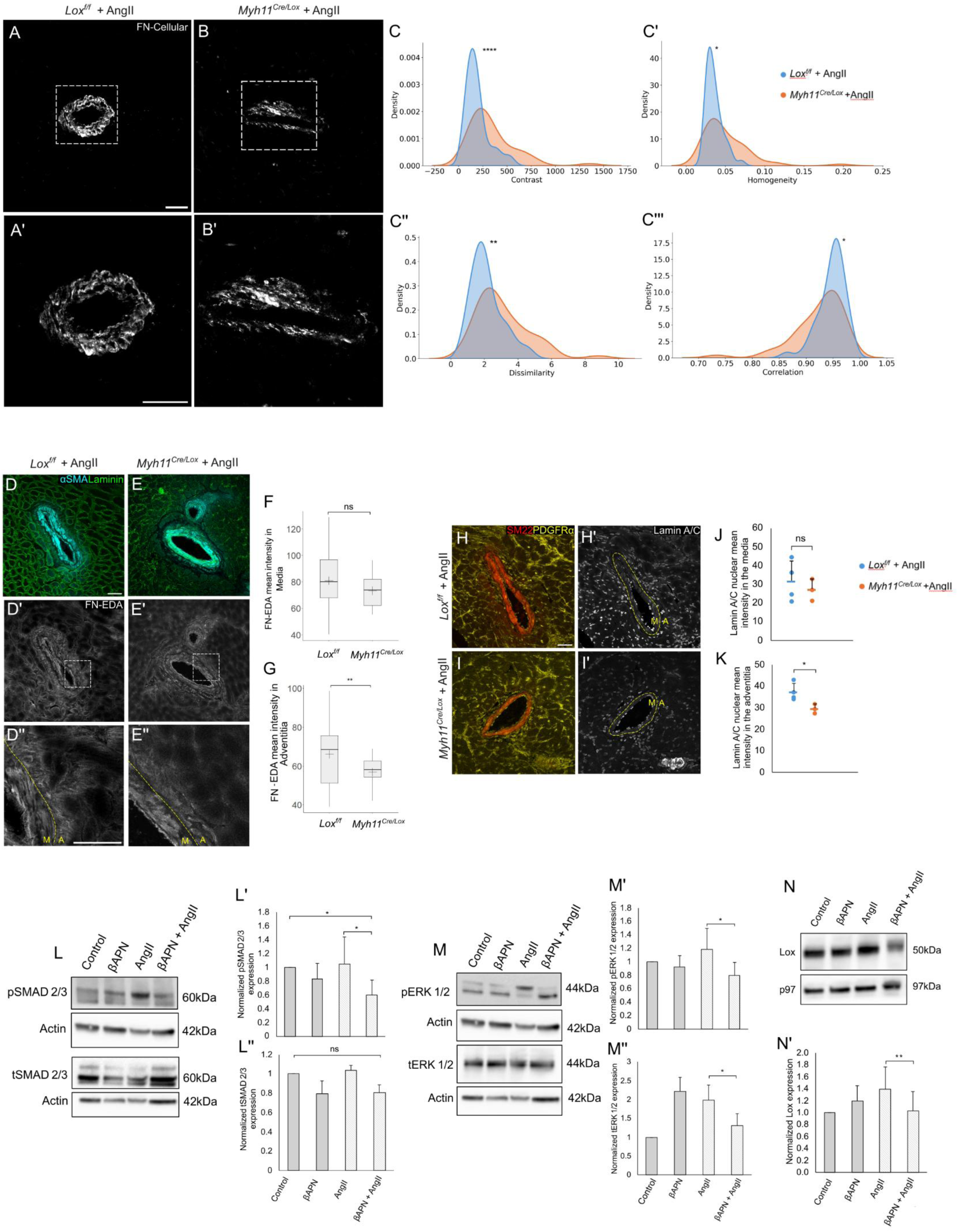
Lox regulates perivascular ECM. Immunostaining for cellular FN in AngII-treated *Lox^f/f^* (A) and *Myh11^Cre/Lox^* (B) mice (Scale bar 50µm). Quantification of Haralick features extracted from cellular FN expression of AngII-treated control *Lox^f/f^* (N=6, n=41) and *Myh11^Cre/Lox^* (N=4, n=48) mice (C). Immunostaining for FN-EDA (grey), αSMA (blue), and Laminin (green) in AngII-treated *Lox^f/f^* (D) and *Myh11^Cre/Lox^* (E) mice’s cardiac vessels (Scale bar 50µm). Quantification of FN-EDA mean intensity in media (F) and adventitia (G) of AngII-treated control *Lox^f/f^* (N=6, n=33) and *Myh11^Cre/Lox^* (N=4, n=12) mice. Immunostaining for Lamin A/C (grey), SM22 (red) and PDGFRα (yellow) in AngII-treated *Lox^f/f^* (H) and *Myh11^Cre/Lox^* (I) mice’s cardiac vessels (Scale bar 50µm). Quantification Lamin A/C expression in media (J) and adventitial regions (K) (Yellow) of AngII-treated *Lox^f/f^* (N=6) and *Myh11^Cre/Lox^* (N=4) mice. Western blot (L,M) and quantification (L’,L’’,M’,M’’) for pSMAD 2/3 and tSMAD 2/3 (N=3), pERK 1/2 and tERK 1/2 (N=5) of HCF cultured on ECM from control and βAPN–treated primary human SMCs in the presence or absence of AngII. Western blot (N) and quantification (N’) for Lox and p97 as control, of HCF cultured on ECM from control and βAPN–treated primary human SMCs in the presence or absence of AngII (N=5). Data are presented as the mean ± SD. Statistical significance is represented by an asterisk as calculated using KS-test, Unpaired t-test, and repeated measures ANOVA with Bonferroni correction. n.s. = not significant. *p < 0.05;**p < 0.01; ****p < 0.0001.

FN-extra domain A (EDA-FN) is a specific subset of cFN that results from alternative splicing^31^. EDA-FN is upregulated following stressful conditions such as wound healing and fibrosis. Notably, EDA-FN induces TGFβ signaling, which in turn induces fibroblast-to-myofibroblast transition ^32^.

Immunostaining specifically for EDA-FN demonstrates that its levels are slightly reduced in the media of *Myh11^Cre/Lox^* mice when compared to that of control. However, in the surrounding adventitial fibroblasts of mutant mice EDA-FN expression is significantly downregulated (Fig. 5D-G).

Mechanotransduction has been associated with hypertensive cardiac remodeling processes. These include alterations to the ECM, changes in the cytoskeleton as well as to the nuclear lamina^33,34^. Having seen the alterations in the medial and adventitial ECM, we wished to test if mechanical signals, signals implicated in fibroblast activation and which are dependent on ECM properties^35^, could be affected in the mutant vessels. One such readout of mechanical signaling is Lamin A/C, a component of the nuclear lamina known to play a role in mechanical signaling to the nucleus^36,37^. Monitoring Lamin A/C expression demonstrates its levels are significantly reduced in adventitia of mutant vessels suggesting that mechanical signaling in the *Lox*-mutant vessels is attenuated (Fig. 5H-K). Collectively, these findings indicate that altered ECM organization following SMC-specific Lox deletion impairs mechanical signal propagation, thereby limiting the fibrotic response in hypertensive hearts.

### SMC-mediated ECM induces fibroblast activation

The above results indicate that outward propagation of the “stress” signal from the vessel wall is attenuated in the hearts of hypertensive *Myh11^Cre/Lox^* mice. Given that Lox is a key enzyme involved in ECM remodeling, these results suggest that signal propagation depends on the mechanical properties of the ECM. Accordingly, ECM produced by SMCs in the presence of AngII could promote the transition of fibroblasts into myofibroblasts.

To directly test this hypothesis, we sought to recapitulate these conditions in vitro using cultured cells and the pan-Lox inhibitor β-aminopropionitrile (βAPN)^38^. To this end, we cultured control and βAPN–treated primary human SMCs^22^ in the presence or absence of AngII. To confirm that the cultured SMCs respond to AngII, we monitored phosphorylated SMAD2/3 (p-SMAD2/3) and phosphorylated ERK1/2 (p-ERK) and their non-phosphorylated versions as their phosphorylation is dependent, in part, on AngII signaling^39^. As expected, AngII treatment increased expression levels of both phosphorylated proteins, indicating these SMCs were responsive to AngII (Fig. S4).

We then cultured SMCs under these conditions for one week to allow deposition of ECM, after which the cells were removed using a decellularization buffer that eliminates cellular material while preserving the ECM^22^. Following extensive washing, primary human cardiac fibroblasts were seeded onto the resulting matrices. After 48 hours, the fibroblasts were lysed and subjected to western blot analysis. To evaluate myofibroblast activation, we assessed levels of p-SMAD2/3 and p-ERK1/2, markers of activated TGFβ and MAPK signaling, respectively, both known to promote fibroblast-to-myofibroblast transition^3^, as well as LOX, which is upregulated in activated fibroblasts^40,41^. We found that cardiac fibroblasts cultured on ECM produced by AngII-treated SMCs exhibited elevated levels of p-SMAD2/3, p-ERK1/2 and LOX compared to fibroblasts cultured on ECM generated under conditions of LOX inhibition (Fig. 5L-N). Together, these results demonstrate a direct link between LOX-modified, SMC-derived ECM and fibroblast activation.

## Discussion

Cardiac fibrosis is a hallmark pathological outcome of chronic hypertension, characterized by excessive ECM deposition that progressively impairs myocardial function^28^. While the distinct roles of various cardiac cell populations in this process are well-documented^3,28,42^, the specific “sentinel” cells responsible for sensing hypertensive stress and orchestrating the subsequent fibrotic cascade have remained unidentified. In this study, we provide evidence that SMCs function as the primary sensors and initiators of the fibrotic response to hypertension, facilitating its propagation from the perivascular regions towards the interstitium. We show this process to be mediated by LOX-dependent ECM remodeling.

### SMCs as Primary Sensors of Hypertensive Stress

A striking finding of our study is the specific decoupling of fibrosis from other hypertensive manifestations. Upon SMC-specific *Lox* deletion, both systemic hypertension and compensatory cardiac hypertrophy remained unchanged following AngII delivery; however, fibroblast activation and interstitial ECM accumulation were significantly attenuated. This suggests that fibrosis is not an inevitable physical consequence of elevated blood pressure, but rather a regulated biological response requiring SMC-mediated signal transduction.

The distinction between SMC and fibroblast roles is further emphasized by the observation that fibroblast-specific *Lox* deletion failed to mitigate the fibrotic phenotype. Despite the high expression of LOX in activated cardiac fibroblasts^25^, and the well established roles of the fibroblasts in the fibrotic response^3^, our data suggest that they act primarily as downstream responders rather than autonomous initiators. This shifts the current paradigm, positioning the SMCs as the critical mediators that sense mechanical stress and subsequently relay activating cues to the surrounding fibroblast population. As such, the SMCs which initiate fibroblast activation, also mediate its propagation into the interstitium.

### LOX-Mediated Fibronectin Scaffolding and Mechanotransduction

Our results suggest that the signaling crosstalk between SMCs and fibroblasts is fundamentally mechanical. We previously established that *LOX* knockdown profoundly alters the composition of the SMC-secreted ECM, specifically targeting fibronectin a highly dynamic glycoprotein that plays a central role in ECM architecture and ECM-mediated signaling, and which is a direct enzymatic substrate of LOX ^10,11,22^. Given that LOX-mediated oxidation is essential for fibronectin fibrillogenesis, which in turn facilitates integrin activation^10,11^ and collagen assembly, our findings imply that SMC-derived LOX is required to maintain the structural integrity of the vascular niche.

In the absence of SMC-derived LOX, the fibronectin scaffold surrounding the media is significantly disrupted. We propose that this scaffold serves as a critical mechanotransduction platform; its perturbation prevents the transmission of mechanical signals necessary for the fibroblast-to-myofibroblast transition and as a result in the *Myh11^Cre/Lox^*mice fibrosis does not spread into the cardiac interstitium. Thus, the SMC-derived ECM acts not only as a structural support, but as a dynamic signaling hub.

Our *in vivo* findings strongly support a mechanotransduction-based model, yet they do not entirely preclude the involvement of LOX-dependent biochemical secretomes. However, our *in vitro* experiments provide definitive evidence that the ECM produced by AngII-stimulated SMCs is independently sufficient to trigger fibroblast activation in a LOX-dependent manner. This confirms that the LOX-modified matrix serves as an active signaling scaffold, converting hypertensive stress into the mechanical cues that drive cardiac fibrosis.

Future studies should aim to further delineate the specific integrin subunits involved in sensing this SMC-derived fibronectin scaffold to identify potential therapeutic targets for uncoupling hypertensive stress from pathological fibrosis.

## Materials and Methods

### Mice

All experiments involving mice conform to the relevant regulatory standards (AAALAC, Technion IACUC protocol #IL083052023H, and to the national animal welfare laws, guidelines, and policies).

*PDGFRα^CreERT^*^2^, *MYH11^CreERT^*^2^ and *Myh11^CreEGFP^*were purchased from The Jackson Laboratory (strain numbers 032770, 019079 and 007742, respectively). Inducible Cre activation was carried out by Intraperitoneal (IP) injections of 50 µl Tamoxifen (Tmx) (Sigma T5648, 20 mg/ml dissolved in 10% EtoH and corn oil) for 5 consecutive days to 3 week-old mice and then 100 µl once a month.

Hypertension induction was carried out via subcutaneous implantation of osmotic minipumps (Alzet Model 2004; 0.25µl/hour, 28 days) loaded with Ang II (AngII; 10mg/1kg/day) to 3 month-old mice. Hearts were harvested 4 weeks following pump implantation.

### Blood pressure

Blood pressure was measured noninvasively in a conscious state by determining the tail blood volume with a volume pressure recording (VPR) sensor and an occlusion tail-cuff (non-invasive tail-cuff CODA System, Kent Scientific, Torrington, CT). Briefly, mice were placed in restrainers on a heating unit and given 15–20 min to acclimatize and reach a steady tail skin temperature (30–35 °C) in a quiet and dark room. Each session consisted of 5 acclimatization measures followed by 15 experimental measurements. Systolic and diastolic blood pressures were calculated from the average of 15 recordings.

### Echocardiography

Echocardiography was performed using a Vevo3100 micro-ultrasound imaging system (VisualSonics, Fujifilm). Maximal left ventricular end-diastolic (LVIDd) and end-systolic (LVIDs) dimensions were measured in short-axis M-mode images at the level of the papillary muscles. Fractional shortening (FS) was calculated as follows:

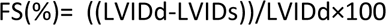

### Histochemistry

Fresh frozen cardiac tissue was serially sectioned at 10 μm intervals. Sections were fixed in 4% Paraformaldehyde (PFA) and Picro Sirius red staining was then performed according to the standard protocol. Images were acquired using Olympus Slideview VS200 slide scanner. FibroTrack software^43^ was used to quantify percentage of interstitial fibrosis by determining the ratio of collagen-stained areas to total area of the heart section.

### Heart tissue immunofluorescence staining

Fresh-frozen 10 µm heart tissue sections were fixed using 4% PFA, then blocked with 5% donkey serum in PBT 0.2%. Sections were incubated over-night at 4 °C or 2h at room temperature with primary antibodies in blocking solution, followed by 2h incubation with secondary antibodies in PBT 0.2% solution.

The following antibodies were used: anti- cellular Fibronectin (Cat# SAB4200784, Sigma, 1:500), anti-SMA (Cat# ab7817, abcam, 1:300), anti-CD31 (DIA310, Dianova, 1:30), anti-GFP (Cat# ab290, Abcam, 1:500), anti-pH3 (Cat# 53348S, Cell signaling, 1:500), anti-PDGFRα (Cat# AF1062, R&D, 1:100), anti-Periostin (Cat# ab14041, Abcam, 1:100), anti- Smoothelin (Cat# 23567-1-AP, Proteintech, 1:300), anti -pFAK (Cat# sc-81493, Santa Cruz, 1:200).

### Fluorescent in situ hybridization (FISH)

FISH was carried out using the Molecular Instruments HCR-RNA-FISH system^44^ according to the manufacturers’ protocol with slight modifications. Briefly, freshly cut frozen tissue sections were fixed using 4% paraformaldehyde for 10 min and then incubated for 1 hour in 70% EtOH. Slides were acetylated using TEA and acetic anhydride, rinsed in DDW, and air dried for 30 min. After pre-hybridizing for 10 min, HCR probes (diluted 1:100) were added to the slides for overnight incubation at 37°C. Slides were washed with Wash- Buffer 5xSSCT gradient then incubated with HCR amplifiers (diluted 1:50) overnight at room temperature. Following this procedure, immunofluorescence was carried out as described above.

### Cell culture

HAOSMCs (Human aortic smooth muscle cell)(ATCC, PCS-100-012) and NHCF-V (Normal human ventricular cardiac fibroblast) (Lonza, Cat# cc-2904) were purchased and cultured according to the supplier’s protocol. Cells were grown and used for experiments up to passage 6. For further maintenance, cells were trypsinized by using Trypsin 0.05% EDTA 0.02% solution (GIBCO, Cat# 25300054) and seeded onto new dishes at density of 5,000 cell/ cm2.

### Preparation of HAOSMC-derived ECM

HAOSMCs were treated with Ang-II 100nM (Alomone Labs SPA-165) or and/or βAPN 500µM in serum free medium (Promocell c-22062) for 5-8 days and allowed to form a tight surface of compact cells and ECM. Cells were removed without affecting the ECM by washing the plate with PBS and then incubated for 1 min with decellularization buffer (1% Triton x-100 and 0.2% ammonium hydroxide). NHCF were then seeded directly onto the ECM (or onto standard tissue culture plates as control) and cultured for 48 hr with full medium (Lonza FGM-3 cc-4526).

### Deep Learning-Based Quantification and Feature Analysis of Fibronectin Images

All analyses were performed in Python 3.10 using Keras-gpu, scikit-learn, OpenCV, and related scientific libraries. Prior to deep learning analysis, images underwent preprocessing steps including blood vessel segmentation and contrast enhancement using Contrast Limited Adaptive Histogram Equalization (CLAHE) with a grid size of 4 and a clip limit of 10. Preprocessed images were then resized to 1024 × 1024 pixels, normalized to a [0, 1] scale, and augmented during training to improve model robustness. A transfer learning approach based on the VGG16 convolutional neural network was employed to classify images as control or mutant, with the model initially trained with frozen weights and subsequently fine-tuned by unfreezing the final convolutional block. Model interpretability was addressed using Gradient-weighted Class Activation Mapping (Grad-CAM) to identify key image regions influencing classification. From these regions, gray-level co-occurrence matrix (GLCM) texture features (dissimilarity, energy, correlation, contrast, and homogeneity) were extracted. Statistical analyses, including principal component analysis (PCA) and linear discriminant analysis (LDA), were performed on the extracted features to assess group differences. All feature data and results were exported to Excel files for final analysis. The final trained model achieved a test accuracy of 88%, with training/validation accuracy and loss curves, as well as a confusion matrix, provided to illustrate model performance.

### Quantitative Analysis of Vascular Tissue Fluorescence

Raw confocal microscopy images were first acquired in czi format and converted into png images following a composite maximum intensity projection step and best fit stretching applied to enhance visualization. Converted images were saved in 8-bit format for subsequent processing. Histological images were then processed using polygonal annotations in the Common Objects in Context (COCO) format to isolate regions of interest corresponding to the adventitia and media, while excluding background (non-vascular tissue and intima). Binary masks were generated for each class (adventitia and media). For each image and annotated region, quantitative metrics including area, integrated intensity, and mean intensity were calculated. All quantitative analyses were performed in Python version 3.12, and the resulting datasets were exported as Excel files for further analysis.

### Imaging

Images were acquired using Olympus BX63 Upright Automated Fluorescence Microscope or the Zeiss LSM 880 laser scanning confocal attached to Axio Examiner Z1 upright microscope (Zeiss, Oberkochen, Baden-W€urttemberg, Germany), with X20 objective and lasers line 405, 488, 561, and 633 nm. Subsequently, all images were processed in a uniform way (average intensity projection, brightness adjustment and pseudo-coloring).

### Cell Area analysis

Fresh-frozen 10 µm heart tissue sections were fixed using 4% paraformaldehyde. Sections were stained with Wheat-germ agglutinin FITC-conjugated (WGA) and images were acquired using Olympus Slideview VS200 slide scanner. Quantification of cross-sectional area (CSA) and cell number per area was performed with Python algorithm.

### Western blot

Heart ventricles from mice or NHCFs lysates harvested from cell culture experiments (Phosphorylation buffer 20mM Tris, 0.3M NaCl, 4mM EDTA, 1% Triton X-100, 1mM Na3VO4, 5mM NaF, 10mM Na4p207) were subjected to SDS-PAGE. Gels were transferred to nitrocellulose membranes, which were then probed with the following antibodies: anti – Actin (Cat# ab179467, Abcam, 1:5000), anti-pERK (Cat# sc-7383, Santa Cruz, 1:500), anti-ERK (Cat# sc-153, Santa Cruz, 1:500), anti – pSMAD 2/3 ( Cat# 8828S, Cell Signaling, 1:1000), anti –SMAD 2/3 ( Cat# 8685S, Cell Signaling, 1:1000), anti-PDGFRα (Cat# AF1062, R&D, 1:1000), anti-Periostin (Cat# ab14041, Abcam, 1:1000), anti-p97 (1:3000; kindly provided by Ariel Stanhill,Open University, IL).

## Acknowledgments

We acknowledge the Imaging Center at Biomedical Core Facility (BCF), Technion – Israel Institute of Technology, for their support throughout the project. We thank M. Holdengreber and M. Gurewitz for their assistance with confocal imaging and slide scanning and Raghd Abu-Sinni for assistance with image analyses. We are grateful to to members of the animal facility for excellent technical assistance. P.H. was supported by grants from the Israel Science Foundation (1126/23), by the Michigan Israel Partnership and by the Rappaport Family Institute. All models and schematics were created with BioRender.com.

## Author Contribution

A.K. conducted the majority of the experiments. A.O. and A.S. carried out image analyses. S.Z.-E. assisted in designing and conducting cell culture experiments. L.C, M.A.S. and R.S. assisted in quantifications and analyses, R.A assisted in *Myh11^CreERT/Lox^* experiment, H.W. and I.K. assisted in designing and analyzing experiment and data. A.K. and P.H. supervised and analyzed the data, planned the study and wrote the manuscript.

## Figure Legends

**Figure S1.**
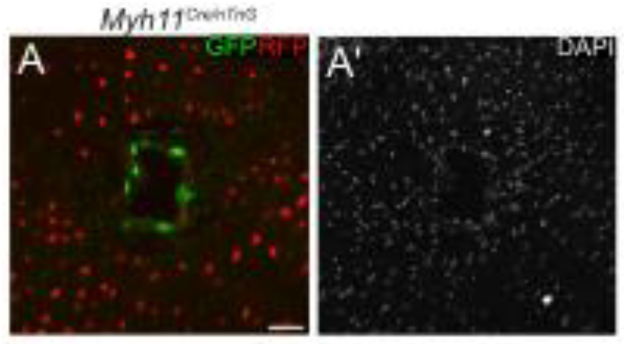
Immunostaining for GFP, RFP, and DAPI in *Myh11^Cre/nTnG^* mice demonstrates Cre activity is specific to SMCs within the media in the cardiac blood vessel (Scale bars 50µm) (A, A’).

**Figure S2.**
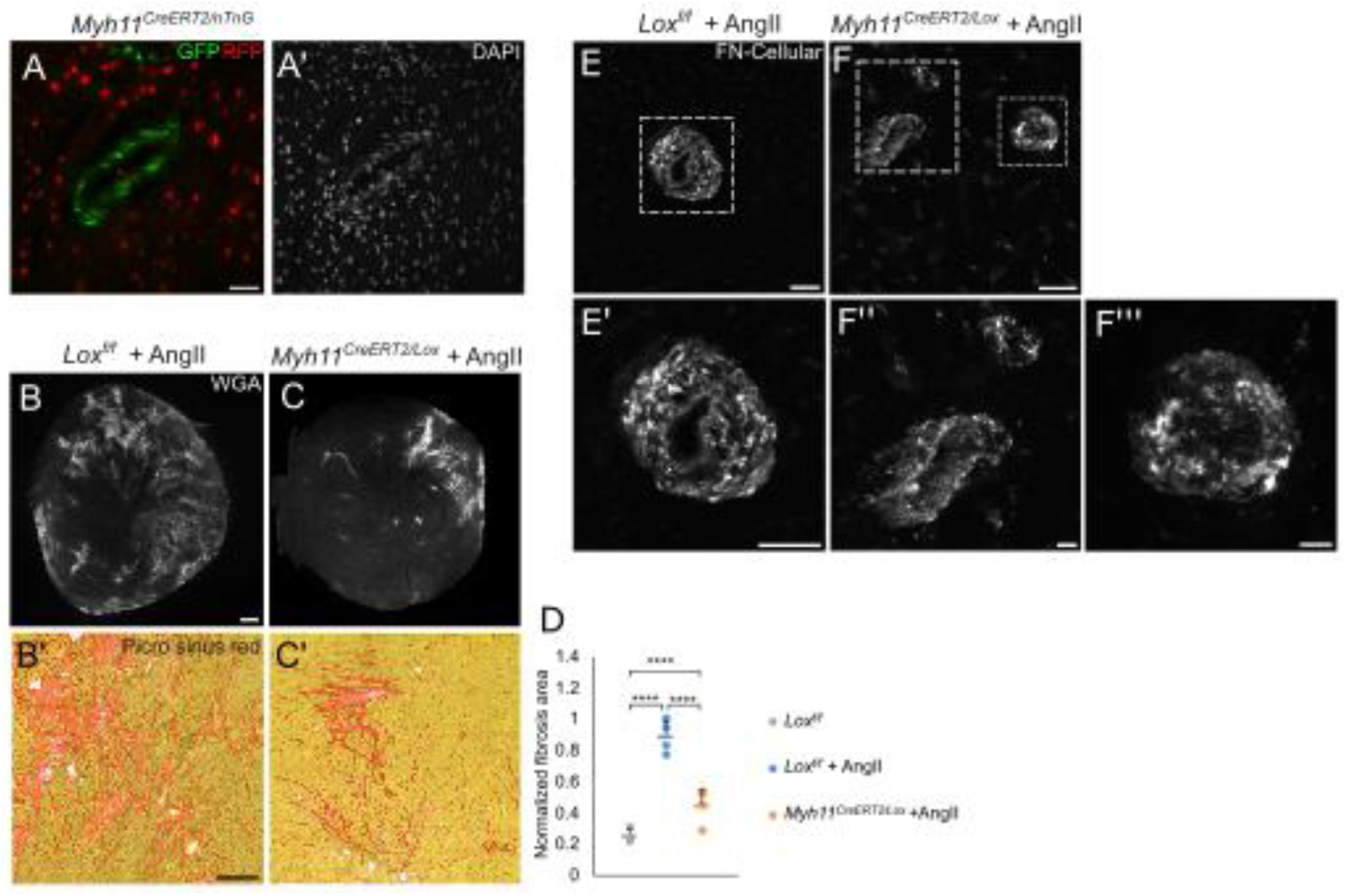
Immunostaining for GFP, RFP, and DAPI in *Myh11^CreERT^*^2^*^/nTnG^* mice demonstrates Cre activity is specific to SMCs within the media in the cardiac blood vessel (Scale bars 50µm) (A). WGA and Sirius red staining (B,C) (Scale bars 500µm, 200µm, respectively) and quantification (D) of baseline *Lox^f/f^* (N=3), hypertensive AngII-treated *Lox^f/f^* (N=4) and *Myh11^CrERTe/Lox^* (N=4) mice. Immunostaining for cellular FN in AngII-treated *Lox^f/f^* (A) (Scale bar 50µm) and *Myh11^CrERT^*^2^*^/Lox^* (B) mice (Upper scale bar 20µm, bottom scale bars 5µm). Data are presented as the mean ± SD. Statistical significance is represented by an asterisk as calculated using one-way ANOVA and post hoc Tukey’s test. n.s. = not significant. ****p < 0.0001.

**Figure S3.**
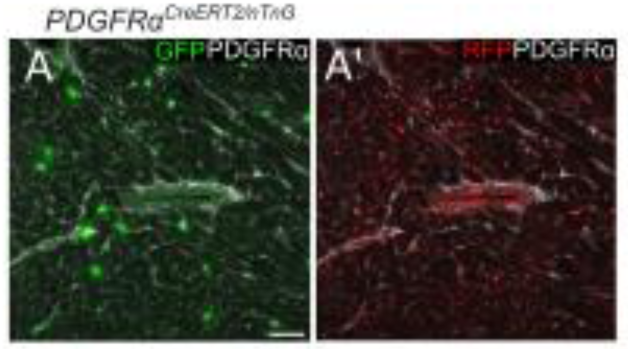
Immunostaining for GFP, RFP, and PDGFRα in *PDGFRα^CreERT^*^2^*^/nTnG^* mice demonstrates Cre activity is specific to PDGFRα-positive fibroblasts within the heart interstitial area (Scale bar 50µm) (A).

**Figure S4.**
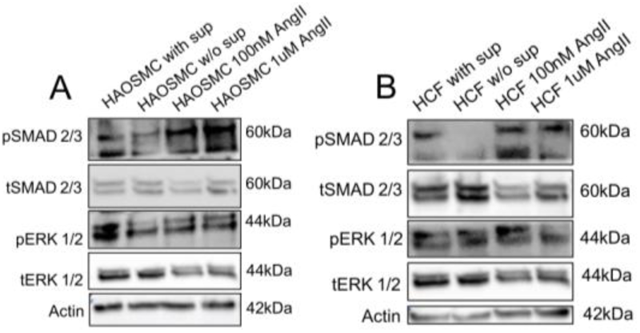
Western blot (A,B) for pSMAD 2/3, tSMAD 2/3, pERK 1/2 and tERK 1/2 of cultured control and βAPN–treated primary human SMCs or HCF in the presence or absence of AngII 100nM or 1µM AngII.

